# Screening and selection of 21 novel microhaplotype markers for ancestry inference in ten Chinese Subpopulations

**DOI:** 10.1101/2021.11.08.467710

**Authors:** Xing Zou, Guanglin He, Jing Liu, Lirong Jiang, Mengge Wang, Pengyu Chen, Yiping Hou, Zheng Wang

## Abstract

Genetic findings suggested that ethnolinguistically diverse populations in China harbor differentiated genetic structure and complex evolutionary admixture history, which provide the genetic basis and theoretical foundation for forensic biogeographical ancestry inference (BGAI). Forensic assays for BGAI among intracontinental eastern Eurasians were previously conducted mainly based on the SNPs or InDels. Microhaplotypes, as a set of closely linked SNPs within 200 base pairs, possess the advantages of both STR and SNP and have great potential in forensic ancestry inference. However, the developed forensic assay based on the ancestry informative microhaplotypes in the BGAI remained to be comprehensively explored, especially in China with enriching genetic diversity. Here, we described a new BGAI panel based on 21 novel identified ancestry informative microhaplotypes that focused on dissected finer-scale ancestry composition of Chinese populations. We initially screened all possible microhaplotypes with high F_st_ values among five East Asian populations and finally employed 21 candidate microhaplotypes in two multiplex SNaPshot assays. Forensic amplification efficiency and statistically/physically phased haplotypes of the 21 microhaplotypes were validated using both SNaPshot and massively parallel sequencing (MPS) platforms. Followingly, we validated the efficiency of these microhaplotypes for BGAI in 764 individuals from ten Chinese populations. Fine-scale ancestry source and ancestry proportion estimated by the principal component analysis (PCA), multidimensional scaling (MDS), phylogenetic tree and model-based STRUCTURE among worldwide populations and East Asians showed that our customized panel could provide a higher discrimination resolution in both continental population stratification and East Asian regional substructure. East Asian populations could be further classified into linguistically/geographically different intracontinental subpopulations (Tibeto-Burman, Tai-Kadai and others). Finally, we obtained a higher estimated accuracy using training and tested datasets in the microhaplotype-based panel than traditional SNP-based panels. Generally, the above results demonstrated that this microhaplotype panel was robust and suitable for forensic BGAI in Chinese populations, which not only provided a high discriminatory power for continental populations but also discriminated East Asians into linguistically restricted subpopulations.

## 1. Introduction

DNA profiling with sets of highly polymorphic autosomal short tandem repeat (STR) has been applied in forensic investigations and has become the mainstream technology in recent years [1–3]. A profile match will provide strong evidence for police investigations. However, the traditional STR method cannot identify the suspects anymore when the profile mismatch. Investigating more valuable information from biological materials found at crime scenes to provide more accurate and reliable guidance for case investigation has now been a new challenge in forensic research. While the biogeographic ancestry information can be inferred through detecting ancestry informative markers (AIMs) of the biological samples, which will provide clues for determining the investigation direction and narrowing the investigation scope. This forensic application was regarded as the forensic biogeographic ancestry inference (BGAI). AIMs are markers that show strong allele frequency differences between populations from different geographic regions or ethnically restricted groups and can be used for determining the probable biogeographic ancestry of an individual [1,2,4,5]. Studies of BGAI have been constructed based on different AIMs, including STR [6], single nucleotide polymorphism (SNP) [7–9] and insertion/deletion polymorphism (InDel) [10–12]. In addition to the traditional AIMs, microhaplotype has become an important part of forensic ancestry inference research in recent years [13,14]. Microhaplotypes, the new genetic markers with two or more closely linked SNPs within 200bp DNA fragment, have the advantages of both STR and SNP and have great application potential in forensic analysis [13–17]. Microhaplotype sets have been developed for forensic ancestry inference, but most of them only focused on inferring ancestry information at the level of intercontinental [13, 17–20]. In recent years, researchers have focused on studies of ancestry inference within intracontinental populations, such as Europe, Southwest Asia [21–23].

China is a large multiethnic country with highly heterogeneous genetic origins and a complex population admixture and evolutionary history. Genetic evidence inferred from the STR, SNP, InDel in the autosomal, X/Y-chromosomes has demonstrated that China with massive ethnolinguistic diversity harbored multiple genetic diversity [24–28]. Recent genetic analyses based on the genome-wide SNP data have revealed the obvious fine-scale population stratifications among the populations from different language families, such as significant genetic differences were identified between Sinitic-speaking Han Chinese and Tungusic/Mongolic/Turkic people in the north [29], Tibeto-Burman people in the southwest [30], and Tai-Kadai, Hmong-Mien, Austronesian and Austroasiatic speakers in the south [31]. Although these aforementioned genetic substructures of Chinese populations have been characterized through SNPs [32], the atlas of the genetic structures of Chinese populations are not fully understood and need further investigation to shed light the fine-scale structures for forensic purposes. To our knowledge, genetic differentiation among intracontinental populations identified via traditionally forensic panels (SNPs, STRs, InDels and others) is limited, even the AIMs included in the Precision ID ancestry panel [33–36].

Therefore, we sought to screened out ancestry informative microhaplotypes for intracontinental population substructure inference. Forty-four markers were screened from five East-Asian populations in the 1000 Genomes Project [37], and then a small sample set (200 individuals) was used to test the efficacy of these loci. Based on the preliminary results, we selected 21 microhaplotypes and constructed two SNaPshot-based panels for typing 764 unrelated individuals from 10 linguistically and geographically different populations. Meanwhile, we designed a massively parallel sequencing (MPS) panel to validate the robustness and reliability of the statistically reconstructed phased haplotypes (PHASE software). Subsequently, populations data from 1000 Genomes Phase 3 were collected and merged with our newly-generated data set to evaluate the efficiency of forensic ancestry inference using these 21 microhaplotype markers. Comprehensive population comparison analyses were performed using different population genetic statistical methods to assess the effectiveness of the 21 loci for substructure discrimination of worldwide continental populations and regional restricted populations. Finally, we performed the ancestry assignment tests using Snipper to evaluate the ancestry assignment capability of this microhaplotype panel.

## 2. Material and methods

### 2.1 Sample preparation

Peripheral blood samples were collected from unrelated individuals after receiving written informed consent. The present study was approved by the Ethics Committee at the Institute of Forensic Medicine, Sichuan University (Approval Number: K2019017), and all the procedures were performed under the standards of the Declaration of Helsinki [38]. A total of 764 samples were collected from ten Chinese populations, including 74 Chengdu Hans, 76 Dujiangyan Tibetans, 77 Muli Tibetans, 78 Xichang Yis, 78 Wuzhong Huis, 63 Zunyi Gelaos, 78 Hainan Lis, 80 Hainan Hans, 73 Ordos Mongolians and 87 Tibet Sherpas. The geographic map of studied populations is shown in **Fig. S1**. All participants shared no biologically close relationships with each other and all of them were required to be aboriginal inhabitants and no marriage with other ethnic groups for at least three generations.

Genomic DNA was extracted using the Purelink Genomic DNA Mini Kit (Thermo Fisher Scientific) and quantified by the NanoDrop-1000 spectrophotometer (Thermo Fisher Scientific) according to the manufacturer’s recommendations. The genomic DNA was diluted to 2 ng/μl and stored at −20◻ until amplification.

### 2.2 Screening and selection of microhaplotypes

The 1000 Genomes Project (https://www.ncbi.nlm.nih.gov/variation/tools/1000genomes/) were used for SNP screening. The following inclusion criteria were used for initial genotyping on the small sample set: (1) candidate SNPs loci with F_st_ > 0.1 among the five East-Asian populations (Southern Han Chinese, China, CHS; Han Chinese in Beijing, China, CHB; Chinese Dai in Xishuangbanna, China, CDX; Kinh in Ho Chi Minh City, Vietnam, KHV; and Japanese in Tokyo, Japan, JPT) were filtered using VCFtools [39]; (2) The physical distance between the two SNPs in one microhaplotype is within 200bp and linkage disequilibrium values between the two SNPs (r2 values) larger than 0.6) [40]; (3) each microhaplotype has at least three haplotypes and all of the haplotypes corresponding minimum frequencies were larger than 0.1; (4) The physical distance among microhaplotypes located on the same chromosome needs to be greater than 20 Mb. According to the above criteria, 44 microhaplotypes were preliminarily screened out. And then the 44 microhaplotypes were further verified and re-screened by testing a small number of samples from ten Chinese populations (20 samples from each population) to find out loci that were suitable for ancestry inference of East-Asian populations. Finally, 21 new microhaplotypes were screened out. The nomenclature of microhaplotypes was in accordance with Kidd’s proposal [41].

### 2.3 Genotyping and Phasing: SNaPshot and PHASE

The PCR primers were designed with the Primer Premier 6.0 [42]. The SBE primers designing was conducted with the SBE primer program [43] and all the SBE primers were tailed at the 5’-end with a poly-GCCTCC(TCCC)n sequence to separate SBE products. All the primers were evaluated for secondary structure and specificity using AutoDimer [44] and NCBI primer blast, respectively. The PCR primers and SBE primers were synthesized by Thermo Fisher Scientific and purified with polyacrylamide gel electrophoresis (PAGE) and HPLC, respectively. The PCR primers and SBE primers of the final 21 microhaplotypes are shown in **Table S1** and **Table S2**.

The PCR reaction was performed in a single PCR multiplex reaction with a total volume of 10 μl, which included 5 μl of QIAGEN Multiplex PCR Master Mix, 1 μl of primer mixture, 1 μl of genomic DNA, and 3 μl of RNase free water. PCR was conducted on the ProFlex 96-Well PCR System (Thermo Fisher Scientific). Thermal cycling conditions consisted of an initial step at 95 ◻ for 15 min, followed by 30 cycles at 94 ◻ for 30 s, 50 - 60 ◻ for 90 s, and 72 ◻ for 30 s, and a final extension at 72 ◻ for 10 min. After the PCR reaction and before the multiplex SBE reactions, in order to remove the remaining primers and nucleotides, we added 2.5 μl of NEB Shrimp Alkaline Phosphatase (SAP) and 0.5 μl of NEB Exonuclease I (EXO I) to 5 μl of PCR products. The mixture was incubated at 37 ◻ for 60 min and then incubated at 80 ◻ for 15 min. Subsequently, the multiplex SBE reactions were performed using the SNaPshot Multiplex kit (Thermo Fisher Scientific) according to the manufacturer’s instructions. 1.5 μl of SNaPshot Multiplex Ready Reaction Mix, 0.5 μl of SBE premixed primers, 1.5 μl of purified PCR product and 1.5 μl of RNase free water were mixed in a total volume of 5 μl. The SBE reactions were conducted according to the following conditions: 25 cycles at 96 ◻ for 10 s, 50 ◻ for 5 s, and 60 ◻ for 30 s. The SBE reaction was performed on the ProFlex 96-Well PCR System. To purify the extension products, 1 μl of SAP was added to 5 μl of extension products and incubated at 37 ◻ for 60 min, followed by incubation at 80 ◻ for 15 min. The purified products were separated and detected by capillary electrophoresis (CE) using the ABI 3130 Genetic Analyzer with POP-7 polymer and GeneScan-120 Liz (Thermo Fisher Scientific). The raw data were analyzed using the GeneMapper ID V3.2 software (Thermo Fisher Scientific). The results of haplotype and corresponding frequencies were estimated by the PHASE version 2.1.1 [45,46]. To assess the sensitivity of the developed SBE panels, a dilution series of template DNA (2 ng, 1ng, 0.5 ng, 0.25 ng, 0.125 ng and 0.0625ng) were amplified and detected in triplicate.

### 2.4 Microhaplotype profiling: massively parallel sequencing (MPS)

MPS technology can obtain haplotype data unambiguously in a single-strand read across the entire locus [13]. To verify the accuracy of the phasing results using PHASE, 119 samples were sequencing by MPS technology. 21 microhaplotype sequence targets were submitted to Thermo Fisher Scientific Ion AmpliSeq Designer (http://www.ampliseq.com) in the form of a BED file (Ampliseq ID: IAD206421). The design type was a single-pool DNA Hotspot design with an amplicon length of 125-375 bp. The Precision ID Library Kit (Thermo Fisher Scientific) was adopted to prepare the DNA library based on the manufacturer’s instructions. After the thermal cycling reaction, we used 2 μl of FuPa Reagent to digest the extra primers in the library products. We added the Switch Solution, diluted barcode adapter mix and DNA ligase successively to add the ligation according to the recommended incubation conditions. After ligation, the Agencourt AMPure XP Reagent (Beckman Coulter) was used to purify the SNP libraries. And then we quantified the obtained libraries using the Ion Library TaqMan Quantitation Kit (Thermo Fisher Scientific) and normalized them to 30 pM. We pooled the libraries in equal volumes for template preparation. The automated template preparation was conducted based on the Ion Chef System (Thermo Fisher Scientific). Subsequently, we used the Ion S5 XL System and Ion 530 chip to sequencing all the microhaplotypes. All raw data were processed with Torrent Suite software V.5.2.2 (Thermo Fisher Scientific), Varianter Plugin V5.2.1.38 (Thermo Fisher Scientific) and Coverage Analysis Plugin V5.2.1.2 (Thermo Fisher Scientific). The BAM files and BAI files were verified using IGV V2.3.97 [47].

### 2.5 Data merging and statistical analysis

The online tool of STRAF [48] was used to evaluate the forensic statistical parameters, including haplotype frequencies, the power of discrimination (PD), probability of matching (PM), power of exclusion (PE), observed heterozygosity (Ho) of the 21 microhaplotypes. The Hardy-Weinberg equilibrium (HWE) and Linkage Disequilibrium (LD) were estimated using the Arlequin v3.5 [49]. Pairwise F_st_ among ten studied populations based on raw genotyped data were also calculated using the Arlequin v3.5. Nei’s standard genetic distances based on the allele frequency spectrum were computed using the PHYLIP v3.6.7 [50]. Principal component analysis (PCA) and multidimensional scaling (MDS) were conducted using the IBM SPSS 23 [51] based on allele frequency and genetic distance, respectively. Phylogenetic trees were established applying the neighbor-joining method using MEGA V.7.0 [52]. Model-based clustering analysis was carried out using the STRUCTURE v2.3.4 [53] with admixture model and correlated frequencies to evaluate the ancestry inference ability of the 21 loci among the ten populations. We ran STRUCTURE from K = 2 to 7. Each K was run 5 times with 100,000 burn-ins and 100,000 Markov Chain Monte Carlo (MCMC) iterations. CLUMPP version 1.1.222 [54] and Distruct version 1.1.23 [55] were used to visualize the STRUCTURE results.

To assess the population genetic relationships among worldwide populations, we collected the population data of 26 populations from the 1000 Genome Project and merged it with our newly-generated population data from ten Chinese populations as the final dataset for the efficiency evaluation of the worldwide and regional ancestry inference and population genetic analyses. Population genetic analyses among the 36 populations were performed using PCA, MDS and neighbor-joining tree, respectively. The population genetic structure was also conducted using the model-based STRUCTURE (K values were set 2 to 8). In addition, ancestry assignments were also performed to evaluate the performance of this inference system with the 21 microhaplotype loci using Snipper 2.5, and 30 of 764 studied individuals were randomly collected as blind trials to estimate their ancestry affiliation.

### 2.6 Quality control

Control DNA 9947A (Thermo Fisher Scientific) and RT-PCR Grade Water (Thermo Fisher Scientific) were used as positive and negative controls for each batch of genotyping, respectively. All experiments were performed at the Forensic Genetics Laboratory of the Institute of Forensic Medicine, Sichuan University, which is an accredited laboratory (ISO 17025), in accordance with quality control measures. The laboratory has also been accredited by the China National Accreditation Service for Conformity Assessment (CNAS).

## 3. Results

### 3.1 Construction of SNaPshot panels

44 candidate microhaplotypes were screened out preliminary from the 1000 Genome dataset. After excluding loci that with small differences in the allele frequencies distribution (F_st_ < 0.05) among the small number of samples (200 samples from ten populations) and eliminating some loci that cannot get stable results via experimental validations, we finally obtained 21 novel microhaplotypes to develop the assay for forensic ancestry inference. The 21 microhaplotypes were distributed on 12 different autosomal chromosomes and the detailed information is listed in **Table 1**. The molecular lengths of the 21 markers ranged from 8 bp (mh03HYP06) to 151 bp (mh08HYP23) with an average of 59 bp, which indicated that these loci might be useful for degraded samples. The electropherogram of the two SNaPshot assays from a reference sample is shown in **Fig. S2**. Serial dilutions of Control DNA 9947A and a reference sample were prepared in triplicate to determine the minimum amount of input DNA. The panel 1 can detect the expected peak when input DNA ≥ 0.0625 ng, and the panel 2 can only detect the expected peak with input DNA not lower than 0.125 ng and the genotype was incomplete at 0.0625 ng.

### 3.2 Validation of the results of genotyping and phasing using MPS

Genotyping by SNaPshot and phasing by PHASE were validated through MPS. 119 samples (11 Chengdu Hans, 16 Dujiangyan Tibetans, 11 Muli Tibetans, 11 Xichang Yis, 13 Wuzhong Huis, 11 Zunyi Gelaos, 11 Hainan Lis, 12 Hainan Hans, 13 Ordos Mongolians and 10 Tibet Sherpas) were used to sequence the 21 microhaplotypes. All the haplotype results estimated by PHASE were in accordance with the sequencing data based on the Ion S5 XL System. These consistent results indicated that genotyping microhaplotypes of SNP-SNP using the method of SNaPshot combined with PHASE is accurate and feasible.

### 3.3 Haplotype frequency distribution and forensic parameters

We successfully genotyped 21 microhaplotypes in 764 individuals from ten Chinese populations and observed 65 different haplotypes in this study. The haplotype frequency distributions of the 21 loci in the investigated populations are shown in **Table S3**. The corresponding haplotype frequencies ranged from 0.0135 (mh09HYP24) to 0.7568 (mh07HYP19) in Chengdu Han, 0.0066 (mh03HYP06) to 0.8026 (mh20HYP42) in Dujiangyan Tibetan, 0.0065 (mh09HYP24) to 0.7857 (mh03HYP06) in Muli Tibetan, 0.0064 (mh09HYP24) to 0.8333 (mh08HYP22) in Xichang Yi, 0.0079 (mh09HYP24) to 0.8413 (mh08HYP22) in Zunyi Gelao, 0.0064 (mh03HYP06) to 0.7308 (mh07HYP19) in Wuzhong Hui, 0.0063 (mh09HYP24) to 0.8333 (mh07HYP19) in Hainan Han, 0.0064 (mh03HYP09) to 0.9103 (mh08HYP22) in Hainan Li, 0.0068 (mh09HYP24) to 0.8493 (mh20HYP42) in Ordos Mongolian, and 0.0115 (mh03HYP09) to 0.9023 (mh11HYP28) in Tibet Sherpa. The haplotype frequency distribution elaborated a large deviation among the ten populations. For example, mh03HYP09 and mh06HYP18 illustrated that distributions of each haplotype in the ten groups were strikingly different (**Fig. S3**). No significant deviation from Hardy-Weinberg disequilibrium and LD was observed among the 21 loci in the ten ethnic groups after Bonferroni correction. The forensic statistical parameters including PM, PD, PIC, Ho, PE and TPI were calculated in each of investigated populations and are presented in **Table S4-S13**, respectively. The Ho values of the 21 loci in the ten populations ranged from 0.0897 (mh07HYP19) in Hainan Li to 0.9310 (mh05HYP14) in Tibet Sherpa with an average value of 0.5216. The cumulative discrimination power (CDP) of the 21 loci spanned from 0.999999999576658 in Hainan Li to 0.99999999999733 in Wuzhong Hui in this study.

### 3.4 Pairwise F_st_ and Nei’s genetic distance

The pairwise F_st_ values among the investigated populations according to genotype data were calculated and listed in **Table S14**. The smallest F_st_ value was observed between Chengdu Han and Zunyi Gelao (0.0044), while the largest F_st_ was found between Tibet Sherpa and Hainan Li (0.0994). Nei’s genetic distances based on the allele frequency distribution were computed to validate and confirm the population’s genetic diversity. As shown in **Table S15**, similar results were obtained. The Nei’s genetic distances varied from 0.0109 (between Chengdu Han and Zunyi Gelao) to 0.2271 (between Hainan Li and Tibet Sherpa).

To explore the worldwide population’s genetic diversity, we also estimated pairwise F_st_ and Nei’s genetic distance among 36 worldwide populations (**Table S16-S17**). The largest Nei’s genetic distance was observed between Mende in Sierra Leone (MSL) from Africa and Tibet Sherpa from Asia (0.4890), while the smallest genetic distance was found between Yoruba in Ibadan (YRI) and Esan in Nigeria (ESN) from Africa (0.0036). Both F_st_ and Nei’s genetic distance revealed that significant genetic distinction existed among intercontinental populations, while close genetic distances exist among intracontinental populations.

### 3.5 Phylogenetic relationship reconstruction and multidimensional scaling analysis

To further investigate genetic relationships among the investigated populations and worldwide populations, neighbor-joining algorithm and multidimensional scaling analysis were conducted on the basis of Nei’s genetic distance matrixes, respectively. In **Fig. 1A**, the neighbor-joining tree of the studied populations presents two main clusters, the Hainan Li and Hainan Han in one branch and the other populations clustered as another main cluster. In the main cluster, Tibeto-Burman-speaking groups (Xichang Yi, Muli Tibetan, Dujiangyan Tibetan and Tibet Sherpa) were found to cluster with each other. The neighbor-joining tree of 36 worldwide populations presents five clusters: East-Asian cluster, South-Asian cluster, African cluster, American cluster and European cluster (**Fig. 2A**). The results of MDS are presented in **Fig. 1B**, Hainan Li was alone in the upper right of the first quadrant, while Hainan Han and Chengdu Han were located close to each other at the lower left. The four Tibeto-Burman-speaking groups of Xichang Yi, Muli Tibetan, Dujiangyan Tibetan and Tibet Sherpa were all in the third quadrant of the coordinate axis. Ordos Mongolian and Zunyi Gelao were in the second and fourth quadrants, respectively. In the MDS of the worldwide populations **(Fig. 2B**), the 36 populations were grouped into three groups: African groups, East-Asian groups as well as European, South-Asian and American groups. African populations were clustered in the lower-left corner, European, South-Asian and American populations were clustered in the middle of the coordinate, and East-Asian populations were distributed on the right side of the Y-axis.

**Fig.1.**
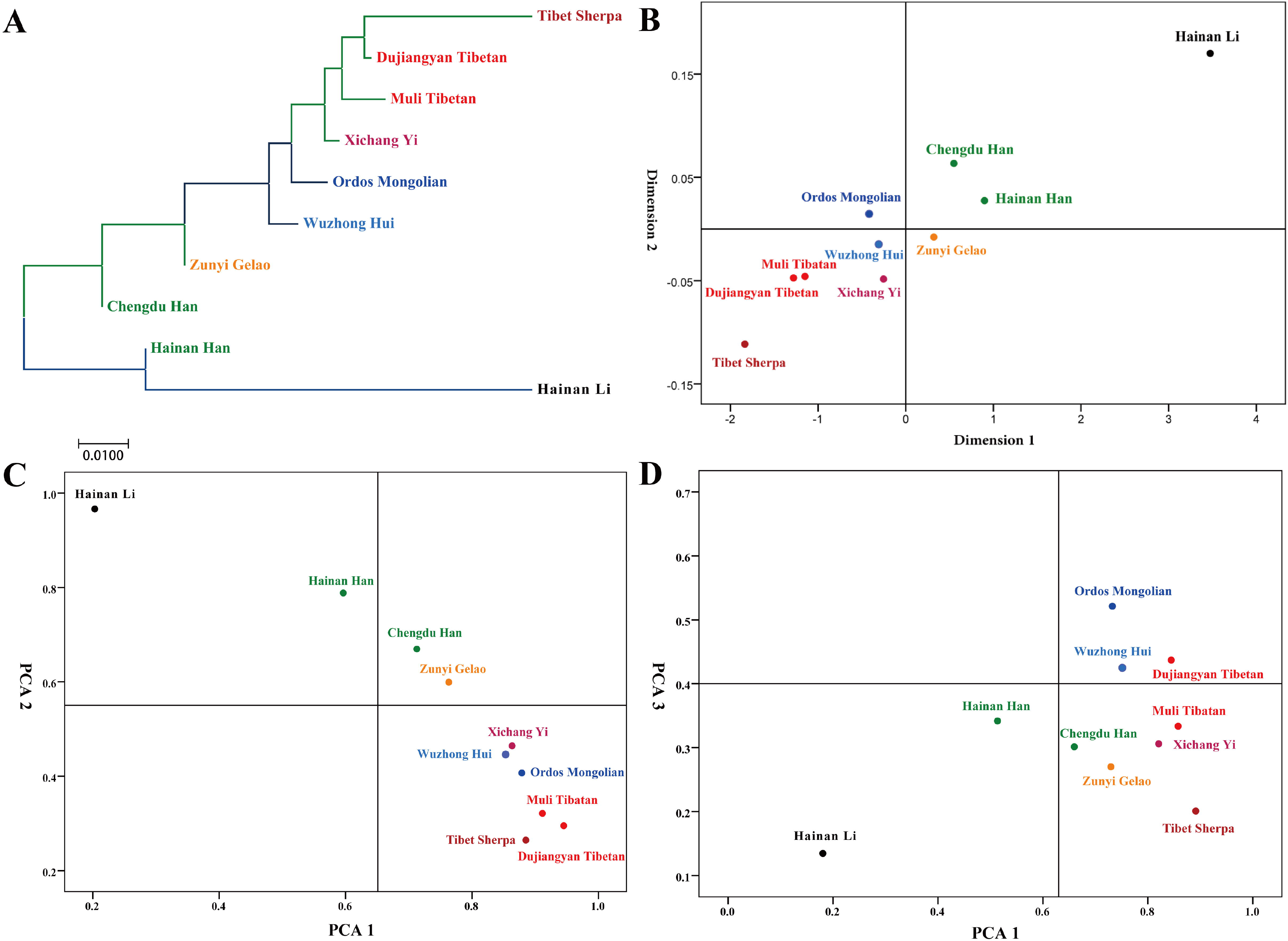
Comprehensive population analyses based on 21 microhaplotypes among the studied populations. (A) A phylogenetic tree among the ten populations based on the Nei’s genetic distances. (B) Multidimensional scaling analysis among the ten populations based on Nei’s genetic distances. (C, D) Principal component analysis based on the first three components among the ten populations. (C) The plot of PC1 and PC2. (D) The plot of PC1 and PC3.

**Fig. 2.**
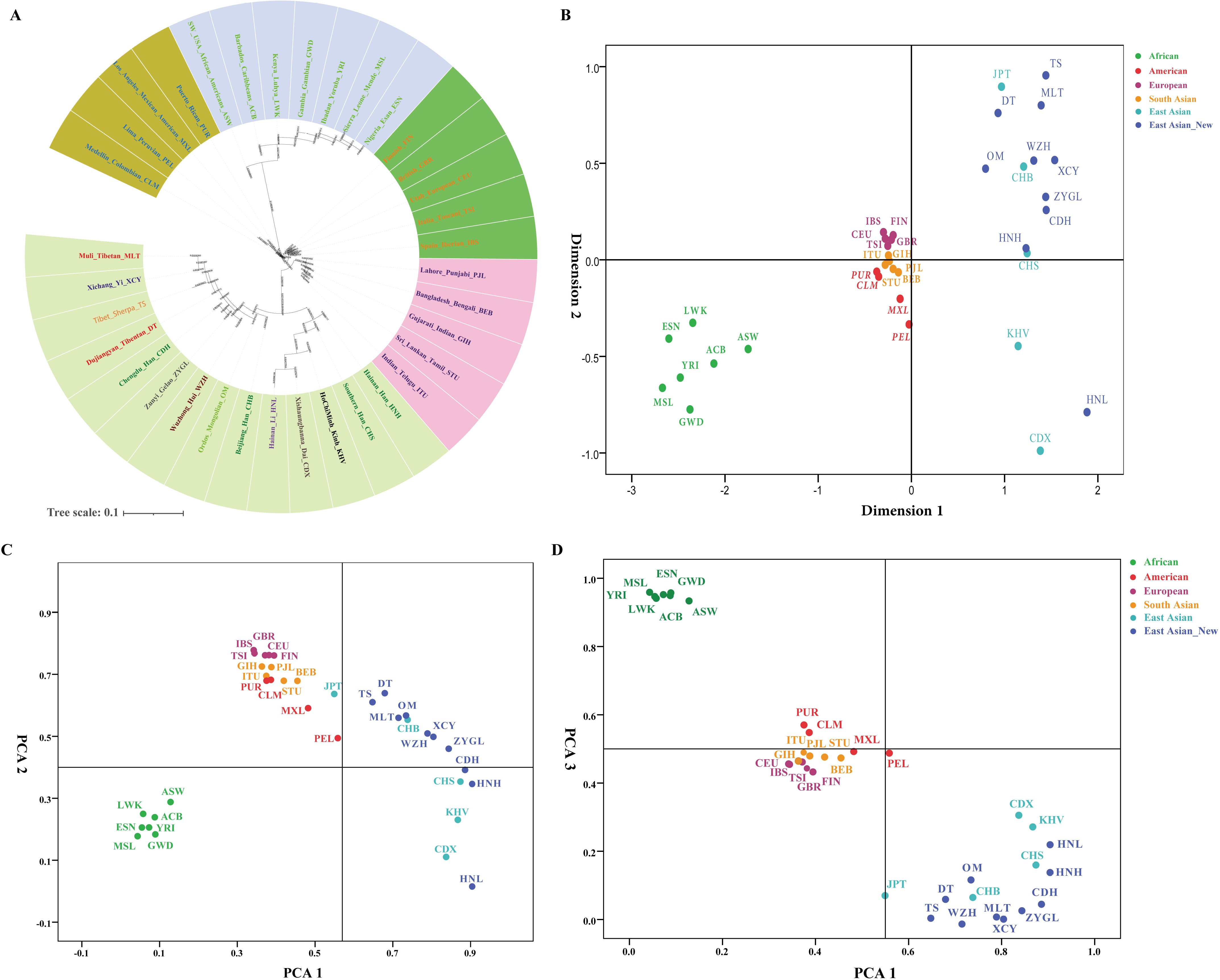
Comprehensive population analyses based on 21 microhaplotypes among 36 worldwide populations. (A) A phylogenetic tree among 36 worldwide populations based on the Nei’s genetic distances. (B) Multidimensional scaling analysis among 36 worldwide populations based on Nei’s genetic distances. (C, D) Principal component analysis based on the first three components among 36 worldwide populations. (C) The plot of PC1 and PC2. (D) The plot of PC1 and PC3.

### 3.6 Principal components analysis

Principal components analysis was conducted based on the allele frequency distribution to assess the population genetic differentiation among the studied and worldwide populations. The first three principal components of the ten studied populations defined 96.5 % of the total genetic variance (PC1: 85.9%, PC2: 8.7%, PC3: 1.9%). The first three principal components presented in **Fig. 1C** and **Fig. 1D** can clearly differentiate Hainan Li and Hainan Han from others. **Fig. 1C** was constructed on the basis of the first two components, and a clear separation between Hainan Li, Hainan Han and other populations was observed. PC1 separated Hainan Li and Hainan Han from the others, and PC2 separated Hainan Li, Hainan Han, Chengdu Han and Zunyi Gelao from the others. In the plot of PC1 and PC3 (**Fig. 1D**), Ordos Mongolian, Wuzhong Hui and Dujiangyan Tibetan were located in the first quadrant. Hainan Li and Hainan Han were separated from each other in the third quadrant. In the PCA plots of 36 worldwide populations (**Fig. 2C** and **Fig. 2D**), the first three principal components can clearly differentiate African populations, Asian populations, as well as European, South Asian and American populations. Although European, South Asian, and American populations clustered in the middle of the coordinate, there were also some genetic differences among each other. PCA results based on allele frequency were similar to the results of MDS on the basis of genetic distance.

### 3.7 Population structures and individual ancestry components

A Bayesian clustering procedure among the ten Chinese populations was conducted using a model-based STRUCTURE algorithm based on genotype data (**Fig. 3**). At K = 2, two ancestral components could be identified. The two principal ancestral components dominate in Hainan Li and Tibet Sherpa, respectively. At K=3, a Hainan-Li-dominant ancestry component can be identified in populations from Hainan and a Tibetan-dominant ancestry component can be observed in Tibeto-Burman-speaking populations. Besides, another ancestry component dominated in Wuzhong Hui and Ordos Mongolian was also found. At K = 4 (the optimal K value), the genetic structure was split by four ancestry components, and Hainan Li, Tibet Sherpa, Wuzhong Hui and Chengdu Han possessed obviously distinct genetic components. With the increase of the K value, the distinction among the sub-populations was separated more clearly. New different ancestral components were continuously separated in the studied populations, while Hainan Li was still composed of only one dominant ancestral component. STRUCTURE was also performed to explore ancestry components among the 36 worldwide populations. **Fig. S4** showed the analysis results of STRUCTURE with K = 2-8. At K= 2, two distinct ancestry components were identified: originating from African and non-African. At K = 4, the 36 populations were grouped into four clusters (African, East-Asian, South-Asian as well as European and American clusters) according to their ancestry components, among which Europeans and Americans were composed of two different ancestry components with different proportions. With the K value increased, the specific ancestry components of different continental populations were shown. At K = 6, African, East-Asian, South-Asian, European and American populations were composed of corresponding specific ancestry components, while two clearly different ancestry components were observed in East-Asian populations. At K = 7, the genetic structure of East-Asians was split into three distinct ancestry components.

**Fig. 3.**
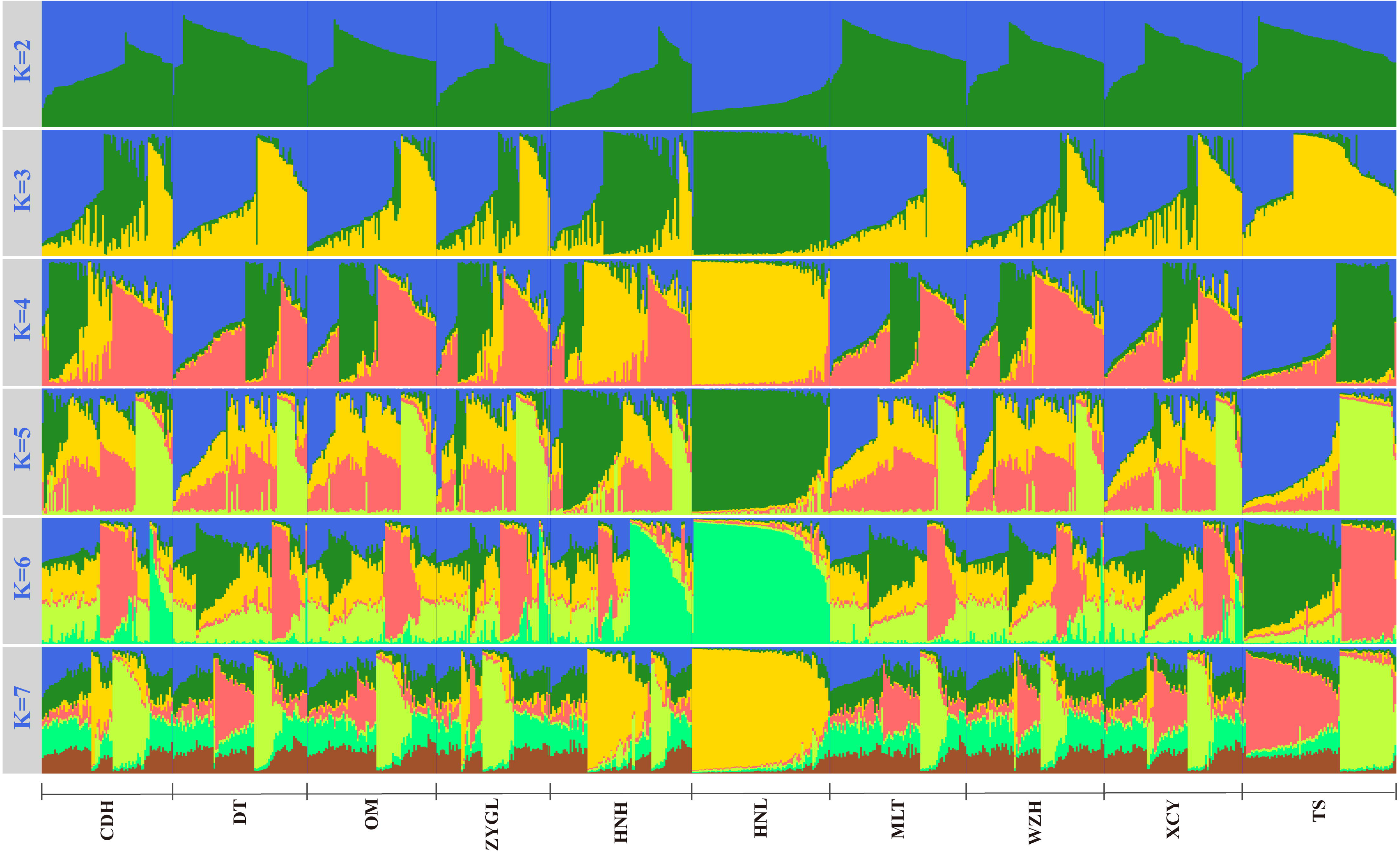
Summary plot of genetic structure among 10 Chinese populations.

Finally, to assess the ancestry assignment performance of the 21 microhaplotypes in Chinese subgroups, we performed an ancestry assignment test using the Snipper (http://mathgene.usc.es/snipper/index.php). A training set (734 individuals from studied populations) and a test set (the remaining 30 individuals from studied populations) were employed. All the test samples were assigned into a most probable population with a list of the resulting likelihood in descending order and the predicted admixtures were also provided. **Table S18** showed the results of the likelihood of all the test samples. In the results of the ancestry prediction test, all the test samples from Hainan Li, Xichang Yi, Zunyi Gelao, Tibet Sherpa and Wuzhong Hui were accurately predicted to the corresponding real population origin. Two individuals from Hainan Han, one from Chengdu Han, one from Dujiangyan Tibetan, two from Muli Tibetan and two from Ordos Mongolian were incorrectly assigned. Although the 8 samples were misclassified as other ethnic groups in neighboring areas or the same ethnic group in different geographical locations, the first three source groups in the list of ancestry probabilities results were all contained their real population sources (**Table S18**).

## 4. Discussion

The ancestry inference of subgroups within a continent is much more difficult than ancestry inference between continents. In order to improve the efficiency of ancestry inference in Chinese subpopulations, we first screened microhaplotype markers based on frequencies distribution among East Asian populations in the 1000 Genomes Project [37]. Owing to the limited East Asian populations in the 1000 Genomes Project, only 44 microhaplotypes were screened out. These loci were further verified and rescreened via 200 samples from different Chinese populations to select suitable loci for ancestry inference of East Asian populations. 21 microhaplotypes with significant population distribution differences in ten Chinese subpopulations were finally screened out. The population data showed that all microhaplotypes had high polymorphisms among the ten subpopulations, with the cumulative discrimination power spanned from 0.999999999576658 (Hainan Li) to 0.99999999999733 (Wuzhong Hui), which indicated that the loci have good application potential in forensic individual identification.

In order to evaluate the ancestry inference efficiency of the 21 loci in Chinese subgroups and to explore the genetic relationship among these populations, the pairwise genetic distances (F_st_ and Nei’s genetic distance) were calculated based on raw genotype data and frequency data, respectively. Moreover, PCA, MDS, phylogenetic analysis and STRUCTURE analysis were conducted to explore the fine-scale population structure and validate the discriminative power of this new-developed panel for regional populations. The results of pairwise genetic distances and neighbor-joining trees showed that the Hainan Li and Tibet Sherpa could be completely separated from other studied populations. This identified population genetic differentiation between Tai-Kadai-speaking Li, Tibeto-Burman-speaking Sherpa and other reference populations was consistent with the recent reconstructed evolutionary and admixture history of these populations based on the genome-wide SNP data. He et al., [31] genotyped over 500,000 SNPs in Hainan Hlai and found these Tai-Kadai people harbored one genetically homogeneous ancestry, which can be used as the representative of the southern Chinese Tai-Kadai ancestry. Further demographic model based on the f-statistics showed that these people shared more ancestry with coastal Austronesian speaking Ami and Atayal in Taiwan island. Recent ancient DNA studies from Fujian and Guangxi also identified two dominant and different Neolithic ancestries (Guangxi ancestry and Fujian ancestry) that played an important role in shaping the patterns of modern genetic diversity of South China and Southeast Asia [56,57]. These genetic analyses showed the southern Chinese ancestry was one dominant ancestry component that needed to be discriminated from others via forensic assay and further provided ancestry inference clues for the forensic investigation. Indeed, our developed microhaplotype-based BGAI panel can successfully differentiate southern Chinese indigenous ancestry from others. Like the unique genetic ancestry identified in Li people, Sherpa-related ancestry was the other typical ancestry component dominant in the highland Tibeto-Burman-speaking populations in the core region of Tibetan Plateau in East Asia [58]. Population admixture history reconstruction based on modern and ancient SNP data showed the genetic differentiation between highland East-Asians and lowland East Asians [30], and these differentiated signatures were also captured by our customized panel. In our population genetic analyses, compared with other groups, Hainan Han had a closer genetic relationship with Hainan Li, and the genetic relationships among the three populations from the Tibetan-Yi corridor (Dujiangyan Tibetan, Muli Tibetan and Xichang Yi) were closer, which suggested that the genetic relationship is also related to the population geographic origin. These results are similar to the results of population genetic analysis based on other genetic markers, such as SNP [34] and InDel [59]. Similar results were also observed in PCA and MDS analysis. In the STRUCTURE analysis, when K was the optimal K value (K = 4), Hainan Li was composed of a specific ancestral component, Dujiangyan Tibetan, Muli Tibetan, Xichang Yi, Wuzhong Hui, Ordos Mongolian and Tibet Sherpa were all composed of three ancestry components in different proportions, while Chengdu Han, Hainan Han and Zunyi Gelao had four distinct ancestral components.

We also explored the genetic relationships among 36 worldwide populations using the 21 loci. The results of MDS showed that the 36 populations were divided into three international groups: African groups, East-Asian groups as well as American, European and South Asian gradient groups. The results of the phylogenetic analysis showed that the African and East Asian populations clustered into two major clusters, while American, European and South-Asian populations first clustered into three branches, and then clustered into a large cluster. PCA results were consistent with the results of MDS and neighbor-joining tree. In the STRUCTURE analysis, with the increase of K value, African, East-Asian, American, European, and South-Asian populations can be completely separated. At the same time, it was also found that Colombians from Medellin (CLM) and Puerto Ricans from Puerto Rico (PUR) from America showed an obvious mixture. In addition to being composed of Native American ancestry, there were also African, European and South-Asian ancestry components in CLM and PUR. Three main East-Asian ancestry components were found in East-Asian populations. Among them, KHV, Hainan Li and CDX were mainly composed of one East-Asian ancestry component. The Tibetan-Yi corridor groups (Xichang Yi, Muli Tibetan, Dujiangyan Tibetan), Ordos Mongolia, Wuzhong Hui and Tibet Sherpa were mainly composed of one other ancestry component, and JPT was made up of another separate ancestral component. The four Han populations (CHB, CHS, Chengdu Han and Hainan Han) and Zunyi Gelao were composed of a mixture of two ancestry components, in which the northern Han (CHB) and the southern Han (CHS, Chengdu Han and Hainan Han) populations had obvious north-south differences. This result is consistent with the research based on high-density SNPs [60]. The abovementioned results suggested that the 21 microhaplotypes not only have good discrimination efficiency in the ancestry inference of intercontinental populations but also in Chinese sub-populations.

For individual ancestry estimation, we used Snipper to analyze all test individuals. We randomly selected 30 samples from studied populations as blind test samples and the remaining samples as the training set. Eight samples were incorrectly predicted to other ethnic groups in neighboring areas or the same ethnic groups in different geographical locations. However, the first three source groups in the results all contained their true source groups. The mismatched results of the above eight samples are mainly related to two factors: the genetic structure of the sample source population and the discrimination efficiency of the genetic markers. When the genetic relationship between the sample source populations of the test set and the reference population of the training set is closer, the prediction bias is more likely to occur, and the requirements for the inference efficiency of the genetic markers is higher. Therefore, it is necessary to screen population-specific and high-resolution AIMs to improve ancestry inference efficiency. In general, the results of Snipper indicated that the 21 microhaplotypes have good potential in ancestry inference of sub-populations in China.

Our study showed that these 21 microhaplotypes are promising ancestry informative markers and have great potential in ancestry inference of Chinese subpopulations and worldwide populations. We also must admit that the efficiency of ancestry inference is limited, and more subpopulations need to be investigated and higher discrimination markers need to be explored in the future. Other limitations in this pilot project focused on the discrimination substructure of Chinese populations via ancestry informative microhaplotypes were the number of included markers, limited SNPs in one microhaplotype, discarding the potentials of other linked InDel or SNP-InDel markers, limited development of specific inference methods and reference databased construction with more linguistically and geographically different populations.

## 5. Conclusion

In this study, we developed and identified 21 novel microhaplotypes as ancestry informative markers, which revealed the powerful potential in ancestry inference. The 21 loci were detected through a SNaPshot and phase workflow and the results were validated using MPS. After the comprehensive analyses of PCA, MDS, neighbor-joining tree and STRUCTURE, results revealed that the 21 loci showed re high-performance for distinguishing populations from East-Asian, African, European, South-Asian and American. Furthermore, the 21 loci were valuable for population stratification in China, which can improve the performance in distinguishing closely resided subpopulations. The 21 microhaplotype-based panel can be used as an effective tool in forensic ancestry inference and population genetics.

## Declaration of competing Interest

The authors declare that they have no conflict of interest.

## Acknowledgement

This study was supported by grants from the National Natural Science Foundation of China (81871532 and 81501635) and the Opening Project of Key Laboratory of Evidence Science (China University of Political Science and Law), Ministry of Education (2021KFKT07). The funders had no role in study design, data collection and analysis, preparation of the manuscript, or decision to submit the manuscript for publication.

---

**Fig. S1** The map showing the geographical location of the investigated Chinese populations.

**Fig. S2** The electropherogram of the two SNaPshot assays from a reference sample.

**Fig. S3** Haplotype frequency distributions of mh03HYP09 and mh06HYP18 in the ten investigated populations in China.

**Fig. S4** Summary plot of genetic structure among 36 worldwide populations.

## Notes

### Competing Interest Statement

The authors have declared no competing interest.

## References

[1] M. Kayser, P. de Knijff, Improving human forensics through advances in genetics, genomics and molecular biology, Nat. Rev. Genet. 12 (2011) 179–192.

[2] M.A. Jobling, P. Gill, Encoded evidence: DNA in forensic analysis, Nat. Rev. Genet. 5 (2004) 739–751.

[3] E. Hagelberg, I.C. Gray, A.J. Jeffreys, Identification of the skeletal remains of a murder victim by DNA analysis, Nature 352 (1991) 427–429.

[4] T.B. Mersha, Mapping asthma-associated variants in admixed populations, Front. Genet. 6 (2015) 292.

[5] M.D. Shriver, R.A. Kittles, Genetic ancestry and the search for personalized genetic histories, Nat. Rev. Genet. 5 (2004) 611–618.

[6] N.A. Rosenberg, J.K. Pritchard, J.L. Weber, H.M. Cann, K.K. Kidd, L.A. Zhivotovsky, et al., Genetic structure of human populations, Science 298 (2002) 2381–2385.

[7] C. Phillips, A. Salas, J.J. Sanchez, M. Fondevila, A. Gomez-Tato, J. Alvarez-Dios, et al., Inferring ancestral origin using a single multiplex assay of ancestry-informative marker SNPs, Forensic Sci. Int. Genet. 1 (2007) 273–280.

[8] Y.L. Wei, L. Wei, L. Zhao, Q.F. Sun, L. Jiang, T. Zhang, et al., A single-tube 27-plex SNP assay for estimating individual ancestry and admixture from three continents, Int. J. Legal Med. 130 (2016) 27–37.

[9] C.X. Li, A.J. Pakstis, L. Jiang, Y.L. Wei, Q.F. Sun, H. Wu, et al., A panel of 74 AISNPs: Improved ancestry inference within Eastern Asia, Forensic Sci. Int. Genet. 23 (2016) 101–110.

[10] N.P. Santos, E.M. Ribeiro-Rodrigues, A.K. Ribeiro-Dos-Santos, R. Pereira, L. Gusmão, A. Amorim, et al., Assessing individual interethnic admixture and population substructure using a 48-insertion-deletion (INSEL) ancestry-informative marker (AIM) panel, Hum. Mutat. 31 (2010) 184–190.

[11] R. Pereira, C. Phillips, N. Pinto, C. Santos, S.E. dos Santos, A. Amorim, et al., Straightforward inference of ancestry and admixture proportions through ancestry-informative insertion deletion multiplexing, PLoS One 7 (2012) e29684.

[12] Q. Lan, C. Shen, X. Jin, Y. Guo, T. Xie, C. Chen, et al., Distinguishing three distinct biogeographic regions with an in-house developed 39-AIM-InDel panel and further admixture proportion estimation for Uyghurs, Electrophoresis 40 (2019) 1525–1534.

[13] K.K. Kidd, A.J. Pakstis, W.C. Speed, R. Lagace, J. Chang, S. Wootton, et al., Current sequencing technology makes microhaplotypes a powerful new type of genetic marker for forensics, Forensic Sci. Int. Genet. 12 (2014) 215–224.

[14] F. Oldoni, K.K. Kidd, D. Podini, Microhaplotypes in forensic genetics, Forensic Sci. Int. Genet. 38 (2019) 54–69.

[15] K.K. Kidd, W.C. Speed, Criteria for selecting microhaplotypes: mixture detection and deconvolution, Investig. Genet. 6 (2015) 1.

[16] J. Zhu M. Lv N. Zhou D. Chen Y. Jiang L. Wang et al., Genotyping polymorphic microhaplotype markers through the Illumina MiSeq platform for forensics, Forensic Sci. Int. Genet. 39 (2019) 1–7.

[17] P. Chen, W. Zhu, F. Tong, Y. Pu, Y. Yu, S. Huang, et al., Identifying novel microhaplotypes for ancestry inference, Int. J. Legal Med.133 (2019) 983–988.

[18] O. Bulbul, A.J. Pakstis, U. Soundararajan, C. Gurkan, J.E. Brissenden, J.M. Roscoe, et al., Ancestry inference of 96 population samples using microhaplotypes, Int. J. Legal Med. 132 (2018) 703–711.

[19] K.K. Kidd, W.C. Speed, A.J. Pakstis, D.S. Podini, R. Lagace, J. Chang, et al., Evaluating 130 microhaplotypes across a global set of 83 populations, Forensic Sci. Int. Genet. 29 (2017) 29–37.

[20] A. Kureshi, J. Li, D. Wen, S. Sun, Z. Yang, L. Zha, Construction and forensic application of 20 highly polymorphic microhaplotypes, R. Soc. Open. Sci. 7 (2020) 191937.

[21] O. Bulbul, G. Filoglu, T. Zorlu, H. Altuncul, A. Freire-Aradas, J. Sochtig, et al., Inference of biogeographical ancestry across central regions of Eurasia, Int. J. Legal Med. 130 (2016) 73–79.

[22] O. Bulbul, W.C. Speed, C. Gurkan, U. Soundararajan, H. Rajeevan, A.J. Pakstis, et al., Improving ancestry distinctions among Southwest Asian populations, Forensic Sci. Int. Genet. 35 (2018) 14–20.

[23] C. Santos, C. Phillips, M. Fondevila, R. Daniel, R.A.H. van Oorschot, E.G. Burchard, et al., Pacifiplex: an ancestry-informative SNP panel centred on Australia and the Pacific region, Forensic Sci. Int. Genet. 20 (2016) 71–80.

[24] H. Yao M. Wang X. Zou Y. Li X. Yang A. Li et al., New insights into the fine-scale history of western-eastern admixture of the northwestern Chinese population in the Hexi Corridor via genome-wide genetic legacy, Mol. Genet. Genomics 296 (2021) 631–651.

[25] M. Wang, G. He, X. Zou, J. Liu, Z. Ye, T. Ming, et al., Genetic insights into the paternal admixture history of Chinese Mongolians via high-resolution customized Y-SNP SNaPshot panels, Forensic Sci. Int. Genet. 54 (2021) 102565.

[26] M. Wang, G. He, S. Gao, F. Jia, X. Zou, J. Liu, et al., Molecular genetic survey and forensic characterization of Chinese Mongolians via the 47 autosomal insertion/deletion marker, Genomics 113 (2021) 2199–2210.

[27] Y. Liu, J. Yang, Y. Li, R. Tang, D. Yuan, Y. Wang, et al., Significant East Asian Affinity of the Sichuan Hui Genomic Structure Suggests the Predominance of the Cultural Diffusion Model in the Genetic Formation Process, Front. Genet. 12 (2021) 626710.

[28] G.L. He, M.G. Wang, Y.X. Li, X. Zou, H.Y. Yeh, R.K. Tang, et al., Fine◻scale north◻to◻south genetic admixture profile in Shaanxi Han Chinese revealed by genome◻wide demographic history reconstruction, J. Syst. Evol. 2021, https://doi.org/10.1111/jse.12715.

[29] G. He, M. Wang, X. Zou, R. Tang, H.Y. Yeh, Z. Wang, et al., Genomic insights into the differentiated population admixture structure and demographic history of North East Asians, bioRxiv. 2021:2021.07.19.452943, doi: https://doi.org/10.1101/2021.07.19.452943.

[30] Y. Liu, M. Wang, P. Chen, Z. Wang, J. Liu, L. Yao, et al., Combined Low-/High-Density Modern and Ancient Genome-Wide Data Document Genomic Admixture History of High-Altitude East Asians, Front. Genet. 12 (2021) 582357.

[31] G. He, Z. Wang, J. Guo, M. Wang, X. Zou, R. Tang, et al., Inferring the population history of Tai-Kadai-speaking people and southernmost Han Chinese on Hainan Island by genome-wide array genotyping, Eur. J. Hum. Genet. 28 (2020) 1111–1123.

[32] O. Libiger, N.J. Schork, A Method for Inferring an Individual’s Genetic Ancestry and Degree of Admixture Associated with Six Major Continental Populations, Front. Genet. 3 (2013) 322.

[33] G. He, Z. Wang, M. Wang, T. Luo, J. Liu, Y. Zhou, et al., Forensic ancestry analysis in two Chinese minority populations using massively parallel sequencing of 165 ancestry-informative SNPs. Electrophoresis 39 (2018) 2732–2742.

[34] Z. Wang, G. He, T. Luo, X. Zhao, J. Liu, M. Wang, et al., Massively parallel sequencing of 165 ancestry informative SNPs in two Chinese Tibetan-Burmese minority ethnicities, Forensic Sci. Int. Genet. 34 (2018) 141–147.

[35] T. Xie, C. Shen, C. Liu, Y. Fang, Y. Guo, Q. Lan, et al., Ancestry inference and admixture component estimations of Chinese Kazak group based on 165 AIM-SNPs via NGS platform, J. Hum. Genet. 65 (2020) 461–468.

[36] G. He, J. Liu, M. Wang, X. Zou, T. Ming, S. Zhu, et al., Massively parallel sequencing of 165 ancestry-informative SNPs and forensic biogeographical ancestry inference in three southern Chinese Sinitic/Tai-Kadai populations, Forensic Sci. Int. Genet. 52 (2021) 102475.

[37] 1000 Genomes Project Consortium, A. Auton, L.D. Brooks, R.M. Durbin, E.P. Garrison, H.M. Kang, et al., A global reference for human genetic variation, Nature 526 (2015) 68–74.

[38] A. Nicogossian, O. Kloiber, B. Stabile, The Revised World Medical Association’s Declaration of Helsinki 2013: enhancing the protection of human research subjects and empowering ethics review committees, World Med. Health Policy 6 (2014) 1–3.

[39] P. Danecek, A. Auton, G. Abecasis, C.A. Albers, E. Banks, M.A. DePristo, et al., The variant call format and VCFtools, Bioinformatics 27 (2011) 2156–2158.

[40] J.C. Barrett, B. Fry, J. Maller, M.J. Daly, Haploview: analysis and visualization of LD and haplotype maps, Bioinformatics 21(2005) 263–265.

[41] K.K. Kidd, Proposed nomenclature for microhaplotypes, Hum. Genomics 10 (2016) 16.

[42] V.K. Singh, A.K. Mangalam, S. Dwivedi, S. Naik, Primer premier: program for design of degenerate primers from a protein sequence, Biotechniques 24 (1998) 318–319.

[43] L. Kaderali, A. Deshpande, J.P. Nolan, P.S. White, Primer-design for multiplexed genotyping, Nucleic Acids Res. 31 (2003) 1796–1802.

[44] P.M. Vallone, J.M. Butler, AutoDimer: a screening tool for primer-dimer and hairpin structures, Biotechniques 37 (2004) 226–231.

[45] M. Stephens, N.J. Smith, P. Donnelly, A new statistical method for haplotype reconstruction from population data, Am. J. Hum. Genet. 68 (2001) 978–989.

[46] M. Stephens, P. Scheet, Accounting for decay of linkage disequilibrium in haplotype inference and missing-data imputation, Am. J. Hum. Genet. 76 (2005) 449–462.

[47] H. Thorvaldsdottir, J.T. Robinson, J.P. Mesirov, Integrative Genomics Viewer (IGV): high-performance genomics data visualization and exploration, Brief. Bioinform. 14 (2013) 178–192.

[48] A. Gouy, M. Zieger, STRAF-A convenient online tool for STR data evaluation in forensic genetics, Forensic Sci. Int. Genet.30 (2017) 148–151.

[49] L. Excoffier, H.E. Lischer, Arlequin suite ver 3.5: a new series of programs to perform population genetics analyses under Linux and Windows, Mol. Ecol. Resour. 10 (2010) 564–567.

[50] M.P. Cummings, PHYLIP (Phylogeny Inference Package), John Wiley & Sons, Inc., 2004.

[51] J. Hansen, Using SPSS for windows and macintosh: analyzing and understanding data, Am. Statistician 59 (1) (2005) 113.

[52] S. Kumar, G. Stecher, K. Tamura, MEGA7: Molecular Evolutionary Genetics Analysis Version 7.0 for Bigger Datasets, Mol. Biol. Evol. 33 (2016) 1870–1874.

[53] G. Evanno, S. Regnaut, J. Goudet, Detecting the number of clusters of individuals using the software STRUCTURE: a simulation study, Mol. Ecol. 14 (2005) 2611–2620.

[54] M. Jakobsson, N.A. Rosenberg, CLUMPP: a cluster matching and permutation program for dealing with label switching and multimodality in analysis of population structure, Bioinformatics 23 (2007) 1801–1806.

[55] N.A. Rosenberg, DISTRUCT: a program for the graphical display of population structure, Mol. Ecol. Notes 4 (2003) 137–138.

[56] T. Wang, W. Wang, G. Xie, Z. Li, X. Fan, Q. Yang, et al., Human population history at the crossroads of East and Southeast Asia since 11,000 years ago, Cell 184 (2021) 3829–3841.21.

[57] M.A. Yang, X. Fan, B. Sun, C. Chen, J. Lang, Y.C. Ko, et al., Ancient DNA indicates human population shifts and admixture in northern and southern China, Science 369 (2020) 282–288.

[58] M. Wang, W. Du, R. Tang, Y. Liu, X. Zou, D. Yuan, et al., Genomic history and forensic characteristics of Sherpa highlanders on the Tibetan Plateau inferred from high-resolution genome-wide InDels and SNPs, bioRxiv 2021:2021.06.23.449553.

[59] X. Zou, G. He, M. Wang, L. Huo, X. Chen, J. Liu, et al., Genetic diversity and phylogenetic structure of four Tibeto-Burman-speaking populations in Tibetan-Yi corridor revealed by insertion/deletion polymorphisms, Mol. Genet. Genomic Med. 8 (2020) e1140.

[60] S. Xu, X. Yin, S. Li, W. Jin, H. Lou, L. Yang, et al., Genomic dissection of population substructure of Han Chinese and its implication in association studies, Am. J. Hum. Genet. 85 (2009) 762–774.

