## Supplementary figures and images for "Screening and selection of 21 novel microhaplotype markers for ancestry inference in ten Chinese Subpopulations"

### Supplementary Figure S1.tif

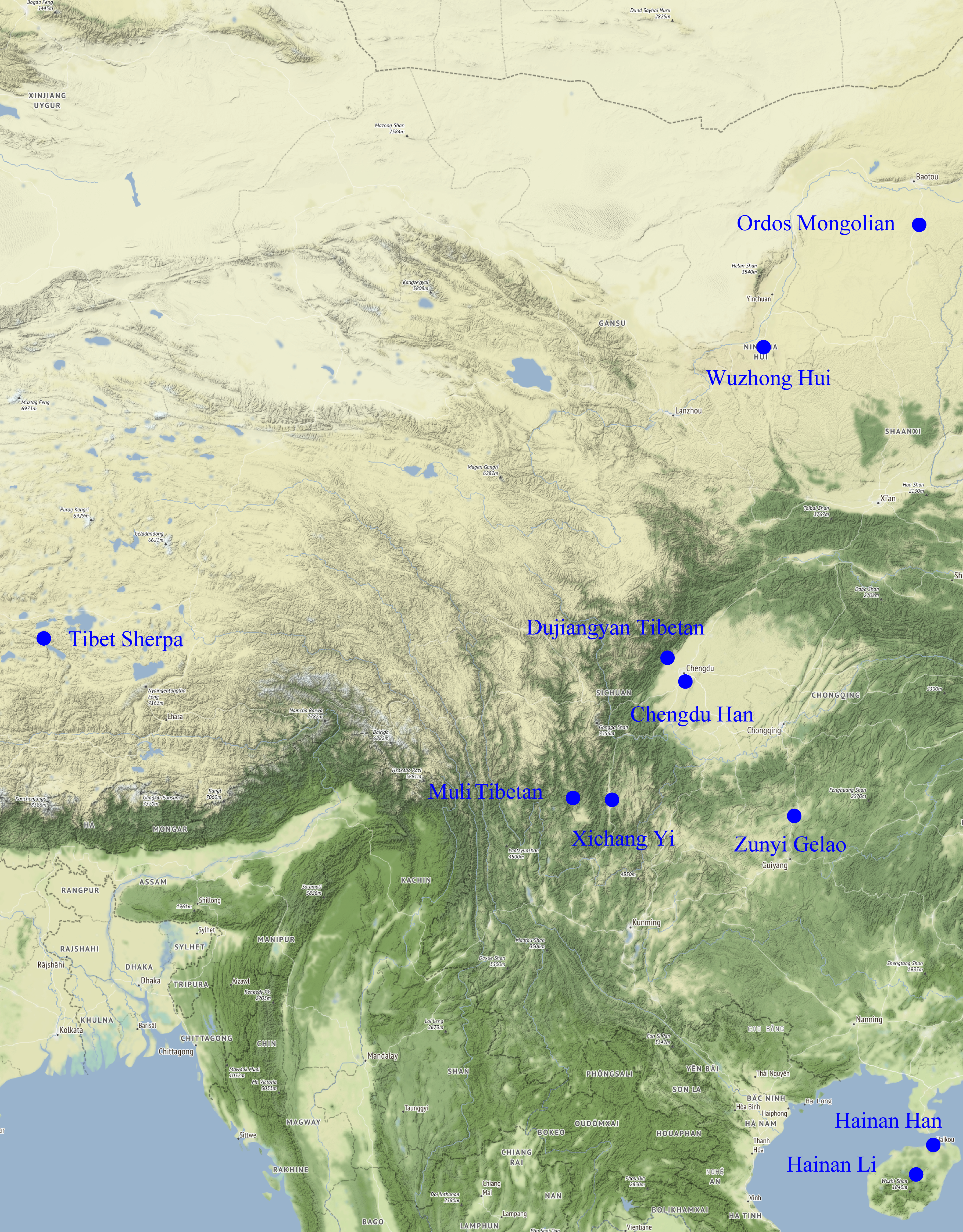

### Supplementary Figure S2.tif

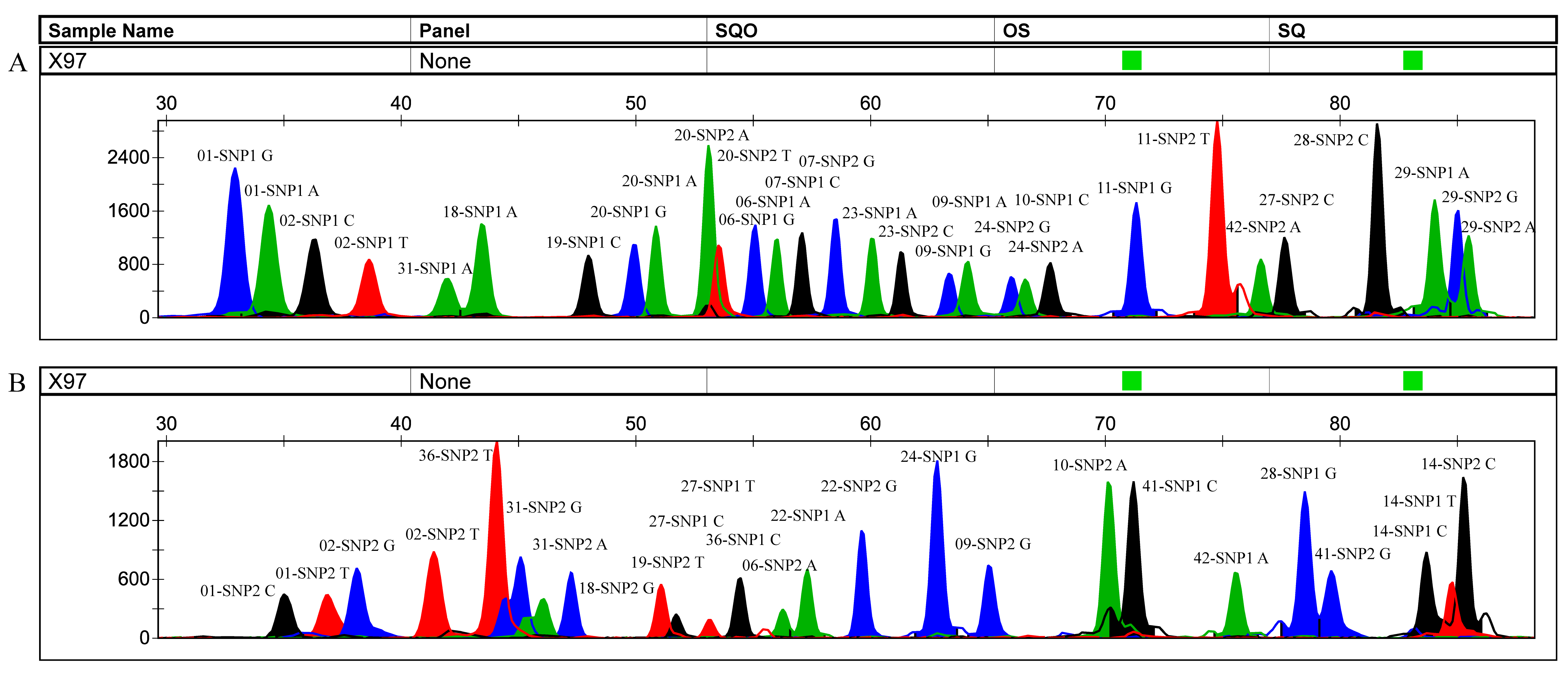

### Supplementary Figure S3.tif

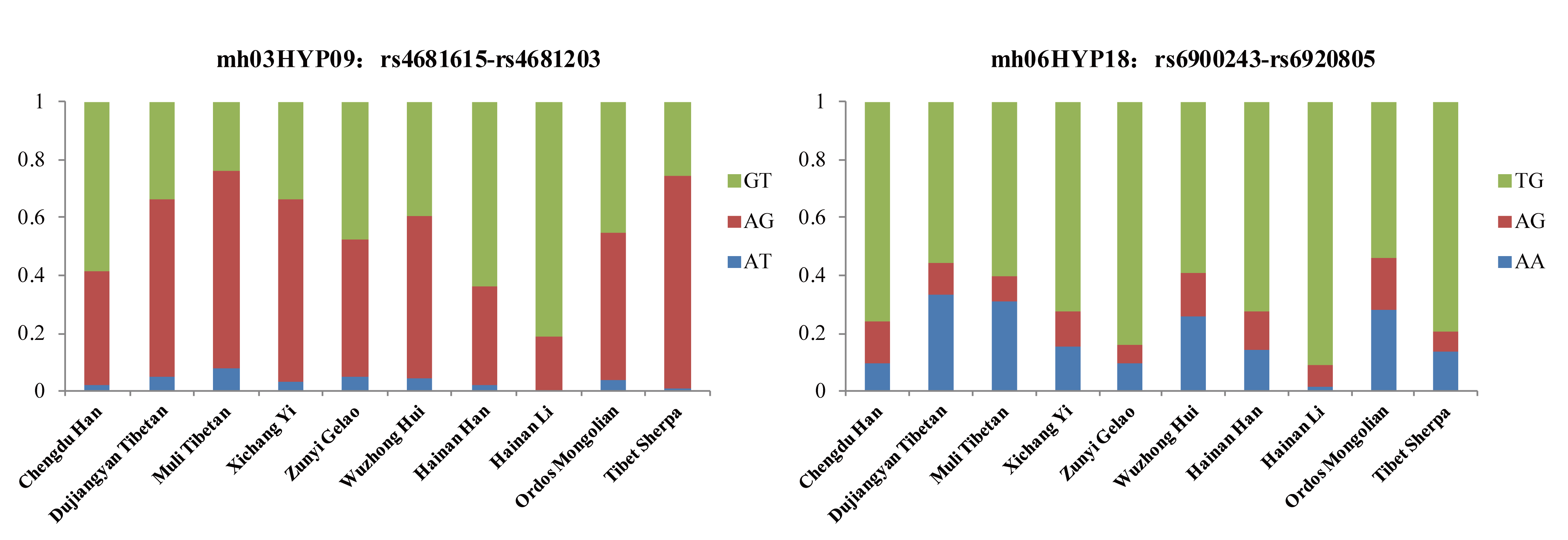

### Supplementary Figure S4.tif

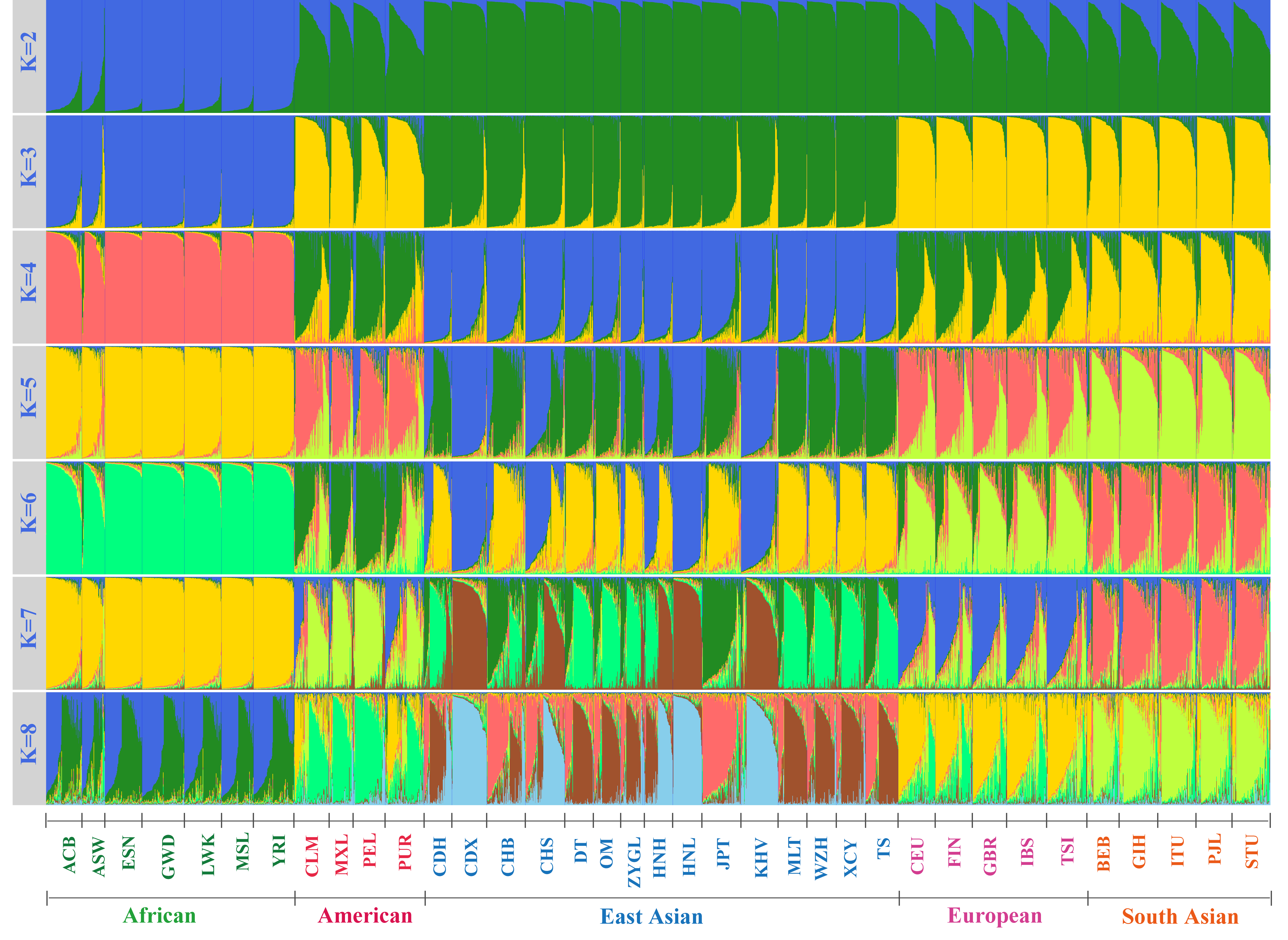
